# Short-term effects of hurricanes Maria and Irma on forest birds of Puerto Rico

**DOI:** 10.1101/578336

**Authors:** John D. Lloyd, Christopher C. Rimmer, José A. Salguero Faría

## Abstract

We compared occupancy in local assemblages of birds in forested areas across Puerto Rico during a winter before (2015) and shortly after (2018) the passage of hurricanes Irma and Maria. Using dynamic community models analyzed within a Bayesian framework, we found significant changes in detectability, with some species becoming more readily detected after the storms and others becoming more difficult to detect during surveys. Changes in occupancy were equally mixed. Five species – mostly granivores and omnivores, but also Black-whiskered Vireo (*Vireo altiloquus*), a migratory insectivore – occupied more sites in 2018 than in 2015. Thirteen species were less common after the hurricanes, including all of the obligate frugivores. Declines in site-occupancy rates were not only more common than increases, but tended to be of greater magnitude. Our results support the general conclusions that bird species respond largely independently to changes in forest structure caused by hurricanes, but that some dietary guilds, notably frugivores, are more sensitive and more likely to show changes in abundance or occupancy following strong storms.

## Introduction

Two significant tropical cyclones crossed near or over Puerto Rico during September 2017. The first, Hurricane Irma, a Category 5 hurricane with wind speeds approaching 300 km h^−1^, passed ∼80 km north of the island on 6 September. The second, Hurricane Maria, made landfall on Puerto Rico on 20 September. A strong Category 4 storm, with maximum sustained winds of ∼250 km h^−1^ at landfall, Maria crossed over the island in a northeasterly direction. Field observations and remotely sensed data revealed that both storms caused significant defoliation of the island’s forests, with Irma causing damage in the northwest corner of the island and Maria defoliating forests and toppling trees across the island [1–3].

Little is known about how the 2017 hurricanes impacted Puerto Rico’s birdlife [4]. Published reports from other islands affected by Maria and Irma are scanty and anecdotal, but suggest that some species were strongly affected [5]. The effects of previous hurricanes on bird populations, especially in the Caribbean, are fairly well documented, however. For most species, the indirect effects of hurricanes via changes in forest structure are far more significant than any direct mortality caused by winds or rain [6]. In the immediate aftermath of damaging storms, birds tend to wander in search of food and shelter. This can take the form of local movements, such as canopy-dwelling species moving into the forest understory [7,8], or larger-scale movements out of heavily damaged habitats or regions [5,9]. Although post-hurricane dispersal of individuals can contribute to changes in the composition and richness of local assemblages, most studies find a relatively small effect of hurricanes in this regard; the species that tend to appear in or vanish from a locale following a storm are typically uncommon or transient members of the assemblage [8–10]. The effect of hurricanes on abundance of individuals is species- and site-specific; in some cases, larger numbers of individuals are encountered after the storm than before, while in others abundance appears lower [7–12]. In general, species that feed on nectar, fruits, and seeds tend to exhibit post-hurricane declines – presumably because availability of these foods is greatly diminished after the storm – whereas numbers of insectivores and omnivores tend to remain stable or increase [6,7,10,12]. However, these patterns are not always apparent [8]. Finally, although local changes in bird assemblages may persist for many years while forest recovery progresses, the population-level effects of hurricanes appear relatively short-lived, with many species recovering to baseline numbers within 6 months [8,11].

Interpreting the biological significance of observed responses by birds to hurricanes is complicated by at least two factors. First, most existing studies are based on surveys carried out over small geographic areas, making it difficult to determine whether changes in observed numbers of individuals reflect changes in population size or simply dispersal away from the study area (for an exception, see [9]). Second, existing research on the effects of hurricanes on birds relies on indices of abundance, whether obtained by capturing individuals in mist-nets or carrying out point-count surveys, that do not address the potentially confounding effect of changes in detectability [8–10]. In particular, significant changes in forest structure and foliage density may make individuals either easier – as in the case of canopy-dwelling species forced into the understory to forage – or harder to detect. Post-storm differences in the number of individuals encountered, then, may reflect either actual changes in abundance (which itself may reflect mortality or emigration) or apparent changes in abundance induced by changes in detectability. Although ad-hoc attempts to address this problem have been employed [8,9], most existing studies of hurricane effects on birds were carried out prior to the development and widespread adoption of analytical methods that control for variation in detectability.

Here, we attempt to address both of these issues by reporting on the results of a geographically extensive survey of forest birds conducted across Puerto Rico before (January - March 2015) and after (January - March 2018) hurricanes Irma and Maria. Analyzing these data within a hierarchical multi-species, multi-season occupancy model [13–16], we sought to document the short-term response of birds to these storms while controlling for the potentially confounding effects of changes in detectability.

## Methods

The surveys reported on here were designed as part of a study of the winter distribution of Bicknell’s Thrush (*Catharus bicknelli*) that was carried out in 2015 [17,18]. Although focused on locating Bicknell’s Thrush, observers recorded all species detected during surveys and as such they offer general insight into the assemblage of forest birds present on Puerto Rico during the winter.

We created a randomized, spatially balanced network of survey locations using a generalized random tessellation stratified (GRTS) scheme. With this approach, we selected 60 1-km^2^ cells from across the island as potential areas in which to conduct surveys. Once we had drawn a sample of cells to survey, we visited each cell and identified 3-5 locations suitable for point-count surveys. Suitability was based on the extent of forest cover – at least 50% of the area in a 50-m radius around each point was forested – and accessibility; all points were along public roads or trails. To maintain independence of counts conducted at different points, we placed each point at least 250 m from its nearest neighbor. Point locations were spatially referenced via GPS and marked with metal tree tags. Due to time limitations – the original survey protocol called for surveys to end in March, prior to any potential pre-migration movement by Bicknell’s Thrush – we only sampled points in 43 of the 60 cells. In total, we sampled 186 points in the 43 cells in both 2015 and 2018. Twenty-five points were surveyed in 2015 but could not be accessed due to storm damage in 2018. To make up for some of these points, we surveyed 16 new points in 2018 within the cells that contained previously surveyed points that had become inaccessible. This yielded 211 points with data from 2015 and 202 points with data from 2018, for a total of 227 points with data from at least 1 year.

Due to the original goal of the surveys, our sample was weighted towards areas more likely to contain habitat for Bicknell’s Thrush based on the winter-habitat model for that species [19] and thus oversamples wet, high-elevation broadleaf forest. Although our survey locations were not a representative sample of all forest types on the island, the sample did include cells at lower elevations in drier forest types. Points surveyed in 2015 ranged in elevation from 0 - 1,297 m, with a median elevation of 705 m (interquartile range = 408 - 825 m). In 2018, surveyed points ranged in elevation from 5 - 1,297 m, with a median elevation of 760 m (interquartile range = 393 - 843 m). Thus, we believe our sampling frame can be described as predominantly forested areas on Puerto Rico that were accessible by roads or trails. The two important sources of potential bias in considering the scope of inference allowed by this sampling frame are 1) the oversampling of high-elevation forests, which will tend to yield low-precision estimates for bird assemblages characteristic of dry, lowland forest and 2) the reliance on trails and roads to access survey points, an unavoidable trade-off given the difficulty of moving through these tropical forests, especially following the hurricanes.

Surveys consisted of standardized 10-minute counts at each point, beginning shortly after sunrise. Each point count was divided into four 2.5-min intervals conducted in immediate succession, with 1-min playbacks of Bicknell’s Thrush vocalizations broadcast before the second and fourth periods. Individuals of all species detected in each count period were recorded in four distance bands (1-10 m, 10-25 m, 25-50 m, >50 m). No counting occurred during the two 1-min playback periods. The use of repeated counts at each point allowed us to model separately the ecological process of occurrence from the observational process of detection [20].

We excluded from analysis all species that were detected on <5% of sites as well as diurnal and nocturnal raptors and waterbirds, species groups that we did not feel were adequately surveyed by our point-count methodology. This left a total of 35 species available for analysis (S1 Table). We converted counts from each point to a binary variable indicating whether or not a species was detected during each of the 4 survey periods. We analyzed presence/absence data using multi-species, multi-season occupancy modeling within a Bayesian framework, treating each point as a site and results from each 2.5-minute interval as replicate counts. Our model treats occupancy as a binary state *z*(*j,t,i*) for each species *i* = 1,2,…,N at site *j* = 1,2,…,J and year t = 1,…,T; *z*(*j,t,i*) = 1 when the species is present and zero otherwise. True occurrence is a latent, unobserved variable, and thus was modeled by a Bernoulli distribution with probability *Ψ*_*j,i,t*_ that species *i* occurs at site *j* and year *t*, specified as *z*(*j,t,i*) ∼ Bern (*Ψ*_*j,i,t*_). The observation model was constructed similarly. Our array of observations *y*(*j,t,k,i*) at site *j*, sampling period *k* = 1,2,…,K, for species *i* at year *t* was assumed also to follow a Bernoulli distribution specified as *y*(*j,t,k,i*) ∼ Bern (*p*_*j,t,k,i*_ * *z*(*j,t,i*)), where *p*_*j,t,k,i*_ was the probability that species *i* at site *j* was detected at period *k* in year *t*. When the species was detected, *y*(*j,t,k,i*) = 1 and is zero otherwise.

We estimated occupancy in 2015 as a species-specific random effect for each species *i*:

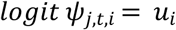

In the 2018, we estimated occupancy as

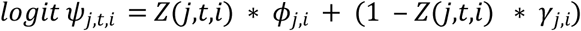

Where *ϕ*_*j,i*_ is the probability that species *i* persisted at site *j* between 2015 and 2018 and *γ*_*j,i*_ is the probability that species *i* colonized site *j* between 2015 and 2018. Although these models can accommodate additional covariates in the process model, we chose to allow occupancy to vary only by species and year. We did so because we did not have any *a priori* hypotheses about factors shaping occupancy, persistence, or colonization rates and because we did not collect any habitat data during surveys. However, we did assume that detectability would vary both among species and between years and, within years, as a function of the date on which the survey was conducted (e.g., because activity patterns vary across the winter) and the time of day (e.g., some species may be more active closer to sunrise and thus more readily detected). As such, we estimated detectability (*p*)as

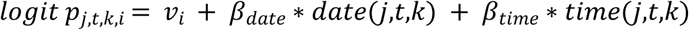

Where *v*_*i*_ was a species-specific random effect, *β*_*date*_ was an additive effect of survey date, and *β*_*time*_ an additive effect of survey time.

In addition to the estimated parameters, we also derived within JAGS values for the change in detectability from 2015 to 2018, change in the number of occupied sites (i.e., the sum of the *Z* matrix in each year), and the rate of change in occupancy, which we calculated as

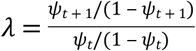

following [21]. For both estimated and derived parameters, we based our inference about statistical significance using the 95% credible interval, which we calculated from the 2.5 and 97.5 percentiles of the posteriors.

We used flat priors for the community level hyper-parameters (i.e., a uniform distribution from 0 to 1 for *ψ, p, ϕ*, and *γ*) and uninformative priors for the survey-level coefficients of detectability (i.e., a random distribution with mean 0 and variance 100) (S1 Appendix). We implemented the analysis using JAGS and the R package ‘jagsUI’ [22,23]. We ran three chains of length 160,000 with a burn-in and adaptation length of 10,000 and thinned the posterior chains by a factor of 10 to reduce autocorrelation, yielding 3,198 posterior samples. These values were set by a process of trial and error, in which we used the shortest chains that still showed evidence of mixing and convergence. We monitored convergence using the Gelman-Rubin diagnostic [24] and used the effective sample size as an indicator of problems with autocorrelation among posterior samples. We also assessed model fit using a Bayesian p-value, which estimates the probability that the simulated data could be more extreme than the observed data [24].

We used general-reference guides to Puerto Rican birds [25,26] to classify species into dietary guilds and followed the nomenclature of [27].

Data used in this analysis are available at https://doi.org/10.6084/m9.figshare.7831424.

## Results

The composition of forest-bird assemblages in 2015 and 2018 was largely the same (S1 Table). Eight species were lost between 2015 and 2018, and 15 were gained. However, all of the species showing turnover between years were uncommon – none occurred at more than 4 sites, and most occurred at only a single site – and most were associated with habitats other than forests (e.g. Royal Tern [*Thalasseus maximus*] or Blue-winged Teal [*Anas discors*]). Much of the turnover, then, reflected incidental detections of transient individuals.

Of the 35 species included in the formal analysis of occupancy, most showed relatively little change in detectability despite substantial changes in forest structure (e.g., defoliation) that might have been expected to influence the probability of detecting birds present during the surveys (Fig 1, S1 Fig). One species, Black-whiskered Vireo (*Vireo altiloquus*), had significantly higher detectability after the hurricanes, although the magnitude of the increase was modest (0.07, 95% CRI = 0.02 - 0.12). Nine species had significantly lower detectability in 2018, with two species – Bananaquit (*Coereba flaveola*) and Scaly-naped Pigeon (*Patagioenas squamosa*) – showing substantially lower detectability in 2018 (−0.27 [95% CRI = −0.23 - −0.31] and −0.32 [95% CRI = −0.24 - −0.41], respectively). Survey date had a mixed but generally weak and negative effect on detectability, with most species showing declines in detectability across the course of the winter (S2 Fig). One notable exception was Black-whiskered Vireo, which had a significantly higher probability of detection later in the season (*β*_*date*_= 0.16, 95% CRI = 0.01 - 0.32), likely a consequence of the large number of migratory individuals that return to the island in February and March to begin breeding. Time of day had weak effects on detectability as well, with only two species having 95% CRI that did not overlap zero (S3 Fig).

**Fig 1.**
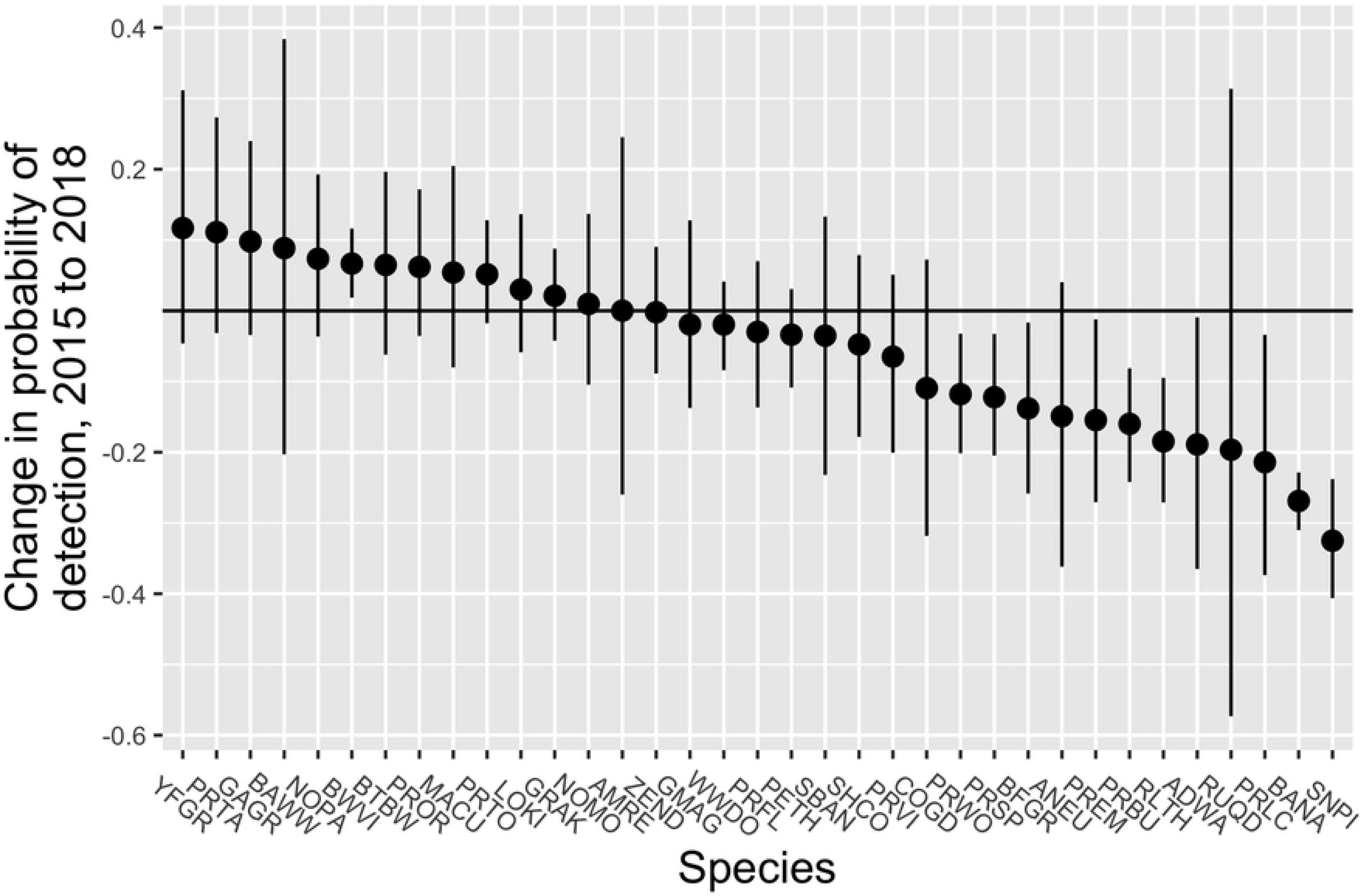
Estimated changes in the probability of detection during winter surveys of forest birds on Puerto Rico before (2015) and after (2018) passage of hurricanes Irma and Maria. Dots show the mean of the posterior samples, and the lines represent the 95% credible interval. For more information on species codes and names, see S1 Table.

Detectability was significantly and positively correlated with occupancy in both 2015 and 2018 (*r* = 0.59 [95% confidence interval = 0.31 - 0.77] and *r* = 0.50 [95% confidence interval = 0.20 - 0.71], respectively; S4 Fig).

Six species showed significant increases in rate of occupancy from 2015 to 2018 (Fig 2, S5 Fig), as evidenced by having 95% CRI that did not overlap with 1: Greater Antillean Grackle (*Quiscalus quiscalus*), Black-faced Grassquit (*Tiaris bicolor*), Northern Mockingbird (*Mimus polyglottos*), Zenaida Dove (*Zenaida asiatica*), Pearly-eyed Thrasher (*Margarops fuscatus*), and Black-whiskered Vireo. Four other species, including two wintering migrants, Black-and-white Warbler (*Mniotilta varia*) and American Redstart (*Setophaga ruticilla*), and two endemic residents, Puerto Rican Emerald (*Chlorostilbon maugaeus*) and Puerto Rican Lizard-Cuckoo (*Coccyzus viellioti*), also showed substantial increases in occupancy, but had very imprecise estimates that overlapped with 1.

**Figure 2.**
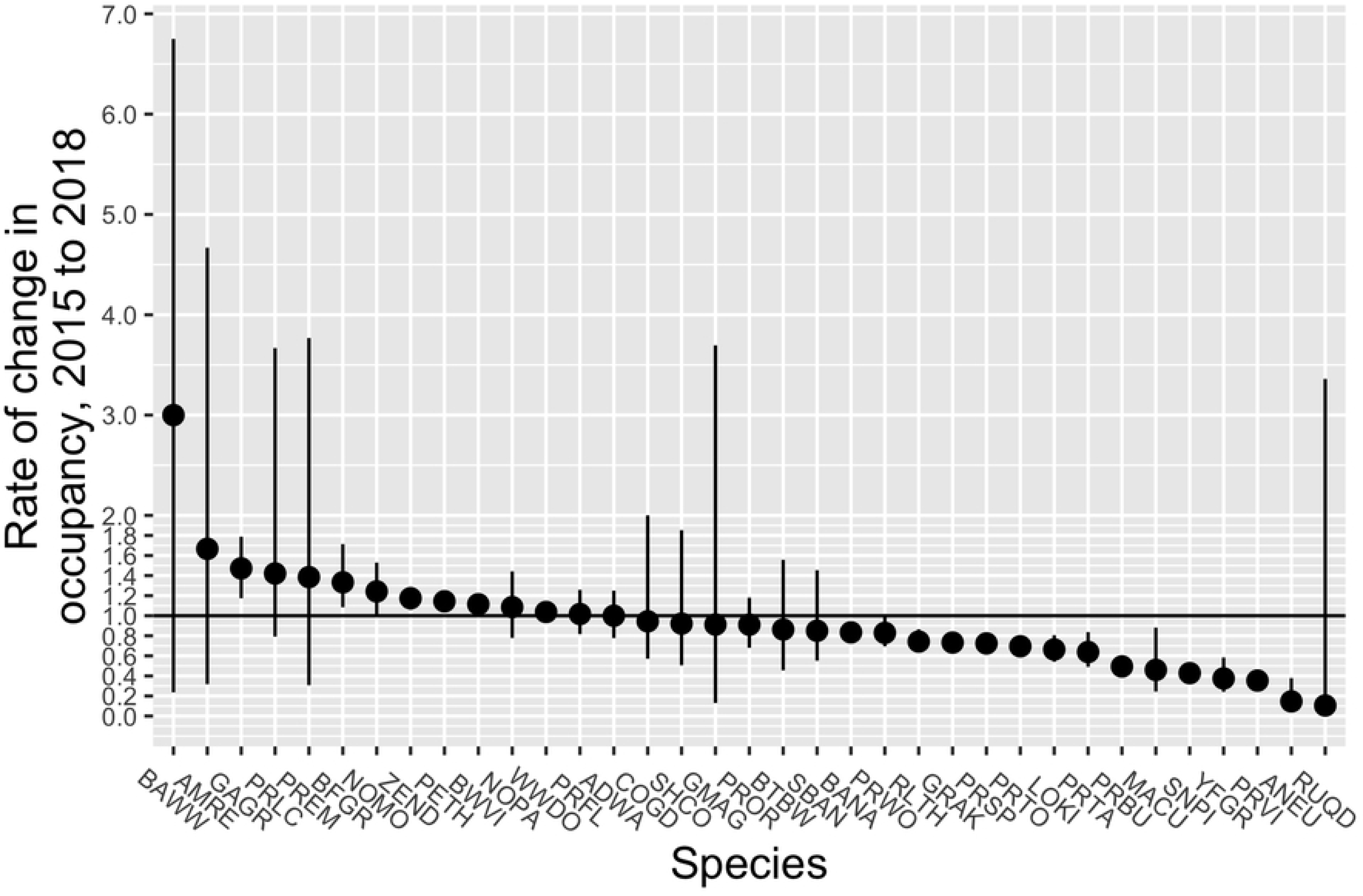
Estimated rate of change in occupancy for forest birds on Puerto Rico before (2015) and after (2018) passage of hurricanes Irma and Maria. Dots show the mean of the posterior samples, and lines represent the 95% credible interval. The solid horizontal line represents no change between years; species falling below the line declined and species above the line increased. For more information on species codes and names, see S1 Table.

Thirteen species had significantly lower probability of occupancy in 2018, including Bananaquit, Red-legged Thrush (*Turdus plumbeus*), Gray Kingbird (*Tyrannus dominicensis*), Puerto Rican Spindalis (*Spindalis portoricensis*), Puerto Rican Tody (*Todus mexicanus*), Loggerhead Kingbird (*Tyrannus caudifasciatus*), Puerto Rican Tanager (*Nesospingus speculiferus*), Puerto Rican Bullfinch (*Loxigilla portoricensis*), Mangrove Cuckoo (*Coccyzus minor*), Scaly-naped Pigeon, Yellow-faced Grassquit (*Tiaris olivaceus*), Puerto Rican Vireo (*Vireo latimeri*), and Antillean Euphonia (*Euphonia musica*). Ruddy Quail-Dove declined substantially (*λ*= 0.49), but occupancy for this species was estimated imprecisely in 2018 – likely because it was encountered on only 3 sites – and thus the 95% CRI around the rate of change was broad (0.03 - 3.33).

The rate of change was larger among declining species than increasing species. Seven species had estimated declines in the rate of occupancy of >50%, whereas none of the species that showed significant increases had estimated gains of >47% (Greater Antillean Grackle:*λ*= 1.47, 95% CRI = 1.17 - 1.79). For example, the largest gain in occupancy probability was by Black-faced Grassquit, which had mean probability of occupancy of 0.24 in 2015 versus 0.32 in 2018 (mean difference = 0.08, 95% CRI = 0.02 - 0.16), whereas all but one of the declining species (Puerto Rican Tanager; mean difference = −0.07, 95% CRI = −0.03 - −0.12) showed decreases of >0.1 in the probability of occupancy and five had declines of >0.2.

The absolute magnitude of change was also greater among declining species than among increasing species (Fig 3). Of the species showing significant increases in occupancy, Black-faced Grassquit gained the most number of sites from 2015 to 2018, 18 (95% CRI = 5 - 36). In contrast, of the declining species, all but Puerto Rican Tanager were estimated to have gone locally extinct at > 25 sites. Antillean Euphonia, Puerto Rican Bullfinch, and Scaly-naped Pigeon all were estimated to have disappeared from >75 sites, with Scaly-naped Pigeon missing from an estimated 102 sites (95% CRI = 89 - 113) in 2018.

**Figure 3.**
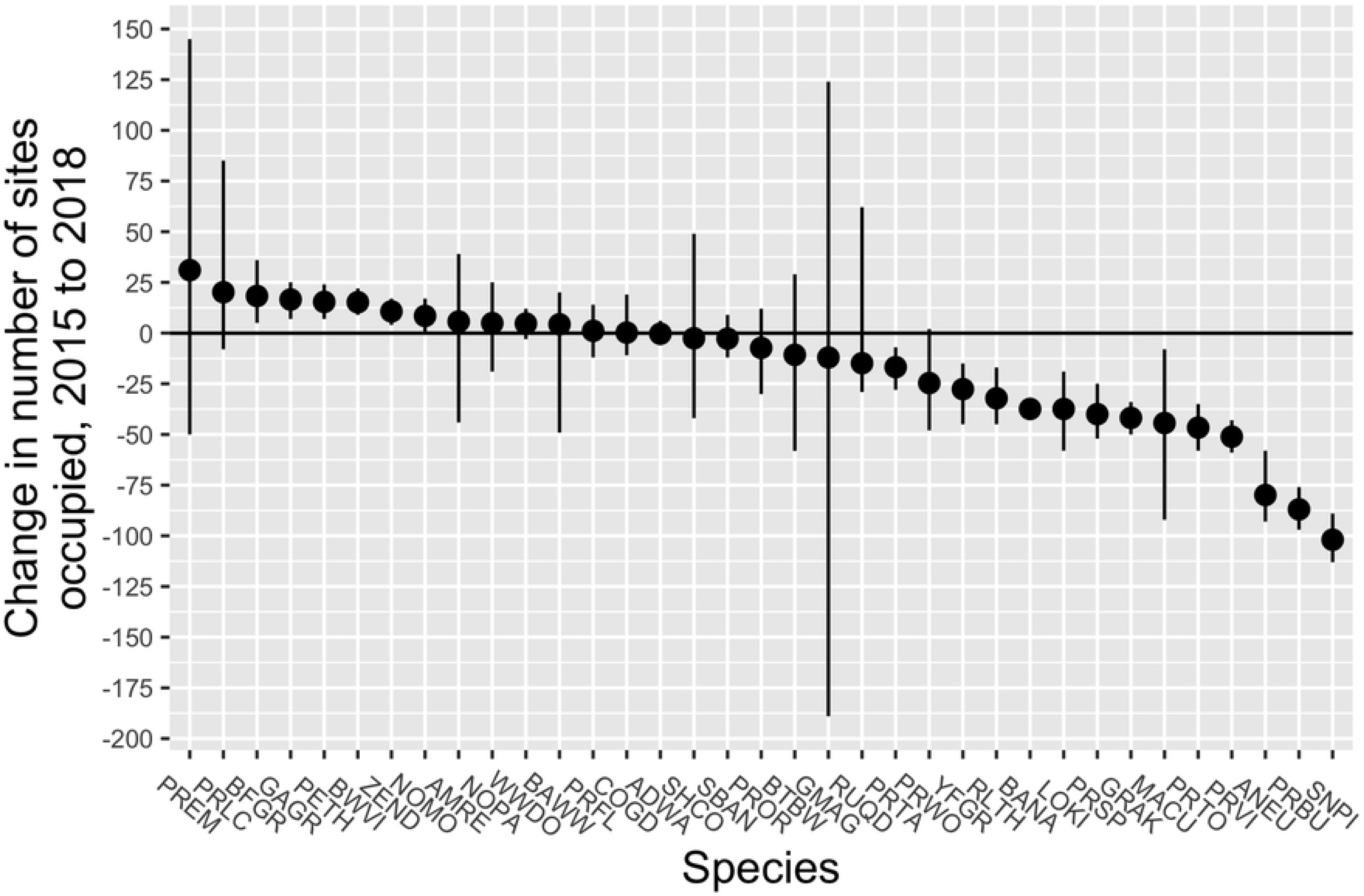
Estimated changes in number of sites occupied by forest birds on Puerto Rico before (2015) and after (2018) passage of hurricanes Irma and Maria. Dots show the mean of the posterior samples, and lines represent the 95% credible interval. The solid horizontal line represents no change between years; species falling below the line declined and species above the line increased. For more information on species codes and names, see S1 Table.

## Discussion

The results presented here draw from one of the few spatially extensive surveys of pre- and post-hurricane bird assemblages in the West Indies, and represent one of the only published reports to explicitly estimate and account for variation in detectability. After accounting for differences in detectability among species and between survey periods, we found that roughly 50% of the 35 species we analyzed showed significant changes in occupancy between our pre-hurricane surveys in 2015 and our post-hurricane surveys in 2018. Far more species appeared to have declined as a result of hurricanes Irma and Maria than to have increased, and the magnitude of the change in occupancy was greater for decreasing species than it was for increasing species.

The first question to address in considering the significance of these findings is whether the changes that we describe reflect actual changes in occupancy and, presumably, abundance [28,29]. Previous research examining the effects of hurricanes on bird populations has acknowledged, although rarely addressed in a systematic fashion, the possibility that changes in indices of abundance, for example from point-count surveys or mist-net captures, reflect changes in detectability but not actual numbers of individuals. Given that hurricanes severe enough to warrant studies of their ecological impact will produce significant changes to vegetation structure in affected areas, which could make individuals much easier or harder to detect, the potentially confounding effects of variation in detectability are a pressing problem.

Our results suggest that detectability varies in a species-specific fashion. Some species became more detectable and others became less so, but relatively few had similar point estimates of detectability between years. Wunderle et al. [9], applying the ad-hoc method of Hutto et al. [30], reached a similar conclusion when comparing detectability of birds on Jamaica before and after Hurricane Gilbert. Thus, not only is bias routine in unadjusted counts, but the direction of the bias is unpredictable and must be addressed individually for each species. This is especially important given the correlation that we observed between occupancy and detectability (see also [31]), which will tend to exaggerate the magnitude of change when uncorrected counts are used. However, changes in detectability and occupancy are not always correlated, and we even found, as in the case of Puerto Rican Lizard-Cuckoo, that the naive estimate of change in occupancy can be of the opposite sign of the estimated change in occupancy due to the magnitude of variation in detectability. We detected this species at 26 (12.3%) and 20 (9.9%) sites in 2015 and 2018, respectively, suggesting a 23% decline. After accounting for imperfect detection (*p* = 0.38 in 2015 v. *p* = 0.15 in 2018), we estimated that Puerto Rican Lizard-Cuckoo in fact occurred at 33 and 47 sites, respectively, representing a rate of increase of >40%. In most cases, unadjusted counts will produce biased – and even worse, unpredictably biased – estimates of change in occupancy. By using repeated point-count surveys to parameterize a dynamic multi-species model that accounts for potential changes in detectability, the estimates of occupancy that we present should reflect real changes in the occurrence of each species.

A second key question is the extent to which the changes in occupancy that we have described can be attributed to the effects of hurricanes Irma and Maria. Our study, like almost every other study of hurricane effects on bird populations, suffers from a lack of replication and lack of control populations. Strictly speaking, this renders speculative any inference about the causal nature of the observed changes. The issue of replication is essentially an unavoidable consequence of the natural experiment afforded by a hurricane; we can subsample extensively, but we are still only measuring the outcome of a single storm occurring in a single place. Including post-hoc control populations is possible but depends entirely on chance, as it requires having had existing monitoring programs in place in areas not affected by the storm [31].

Although we acknowledge the possibility that changes in occupancy for some or all species in our analysis might be caused by factors other than the hurricanes, three patterns suggest that at least some of the described changes are causally linked to the hurricanes. First, we found that declines in occupancy were widespread and strong among frugivores, especially so for Scaly-naped Pigeon, Puerto Rican Bullfinch, and Antillean Euphonia. Both anecdotal and empirical evidence suggest that hurricanes often eliminate standing fruit crops [32,33] and previous research has consistently reported declines in indices of abundance among frugivorous birds – including those included in our analysis – and bats following the passage of a hurricane [9,10,12,34,35]. Thus, the observed declines in occupancy for the five obligate frugivores are consistent with the expected indirect effects of a hurricane.

Second, of the species that increased in occupancy following the storm, all but one – Black-whiskered Vireo – are habitat generalists that will use open, non-forested environments, including human-dominated landscapes, and forest edges [25]. Not only are these types of environments less prone to hurricane damage, thus reducing the probability of local extinctions, but the creation of gaps and extension of forest edge due to wind damage may actually have increased the extent of habitat available for these species. Indeed, post-hurricane increases in open-country species not normally found in forests, or detected only in low numbers, is a common finding [8,32,36]. This pattern, too, is consistent with a causal effect of hurricanes Irma and Maria.

Finally, we observed a substantial decline in occupancy among Bananaquits, a species often considered sensitive to hurricanes due to its reliance on nectar as a food source. Indeed, many studies have documented declines in Bananaquits after hurricanes [10–12,37,38] or other disturbances that reduce floral abundance (e.g., [39]).

Many of the other observed changes in occupancy are less clearly linked to the effects of hurricanes Irma and Maria, both because we lack an understanding of the likely mechanism by which hurricane damage may impact other species or guilds and because other studies report conflicting results. For example, whereas our results and those of Waide’s [11] show post-hurricane increases in Pearly-eyed Thrasher, a common omnivore, Askins and Ewert [12] reported significant declines in the species on St. John, US Virgin Islands, following Hurricane Hugo. Likewise, we found that Puerto Rican Tody had lower occupancy following the hurricanes, but Waide [11] reported that this species increased in numbers after Hurricane Hugo. We observed a large decline in occupancy among Yellow-faced Grassquits (*Tiaris olivaceus*), yet this species is a common resident of roadsides and open areas [40,41], environments not obviously sensitive to effects of hurricanes.

One of the challenges in interpreting results of pre- and post-hurricane surveys, and in making sense of the apparently conflicting results evident in the literature, is that how species respond to hurricanes likely reflects some combination of inherent, trait-based sensitivity to disturbance and the degree of impact that storms have on critical resources. Although species that depend on resources easily damaged by hurricanes – fruit or nectar, for example – may be inherently more vulnerable to effects of hurricanes, these effects may only emerge if storm damage is severe. Conflicting results regarding the apparent sensitivity of different species can thus emerge simply because most studies sample relatively small areas that may not incorporate the full gradient of disturbance intensity. Indeed, studies that have incorporated geographically extensive surveys tend to show this pattern clearly: even within a species, patterns of response to the same storm vary from location to location, with species tending to decline in heavily damaged areas and increase or remain stable in areas where damage was relatively light [9,42]. Future research would therefore benefit from sampling bird populations across geographically extensive areas and from monitoring in these same locations the resources – for example food or suitable nest sites – that putatively underlie population-level responses to storm damage. Doing so would provide a more complete picture of how populations respond while at the same time facilitating identification of causal links between hurricanes and demographic change. Because hurricanes are infrequent and unpredictable in their timing, however, this would necessitate implementing and maintaining large-scale ecological monitoring programs.

For the changes in occupancy that seem plausibly linked to the damage caused by hurricanes Irma and Maria, a final question is whether they reflect changes in population growth rate or temporary emigration. Although significant demographic impacts are possible, especially for species with small population sizes [5,43], some of the apparent changes in abundance or occupancy may reflect temporary emigration from areas where vegetation has been heavily damaged. For example, we observed a substantial decline in Scaly-naped Pigeons, yet in previous hurricanes this species has responded by emigrating from areas with extensive damage to areas where forests remained relatively intact [42]. Given their prodigious dispersal capacity, Scaly-naped Pigeons may be somewhat unique in their resilience to widespread storm damage. A species’ capacity to respond to hurricane damage via emigration likely depends on both the extent of damage and the dispersal ability of that species; emigration may not be an effective strategy for species with relatively poor dispersal abilities facing geographically extensive damage. If Puerto Rico’s endemic birds, like other island endemics [44–46], possess relatively weak dispersal abilities, then observed declines in Puerto Rican Bullfinch, Puerto Rican Vireo, Puerto Rican Tody, Puerto Rican Spindalis, and Puerto Rican Tanager more likely reflect population declines than movement out of our study area. An additional complication is that many studies of hurricane effects – ours included – sample populations over short time scales, which may exaggerate the significance of hurricanes as a driver of population change, especially if emigration is the primary initial response. The few post-hurricane studies that have monitored bird populations over longer time periods show that most species are resilient in the face of hurricane damage [8,11,42].

In conclusion, our results add to the body of empirical findings regarding the effects of hurricanes on birds, and support the general conclusion that bird species respond largely independently and presumably in response to changes in forest structure caused by these storms. Although some species appeared to benefit from the changes caused by hurricanes Irma and Maria, we found that negative effects, in the form of declines in site-occupancy rates, were stronger and more widespread. Further, we found that detectability was a potentially significant source of bias in comparisons of pre- and post-storm occupancy. That detectability and occupancy were correlated suggests that uncorrected comparisons of occupancy are likely to exaggerate the effects of storms.

## Acknowledgements

We are grateful for the excellent field assistance provided by Alcides Morales, Carina Roig, Julio Salgado, Pedro Santana and Fabio Tarazona. We thank the Sociedad Ornitológica Puertorriqueña, Inc. for providing invaluable logistic advice.

## Supporting information

**S1 Figure. Probability of detection during winter surveys of forest birds on Puerto Rico before (2015) and after (2018) passage of hurricanes Irma and Maria.** Dots show the mean of the posterior samples, and lines represent the 95% credible interval. For more information on species codes and names, see S1 Table.

**S2 Figure. Value of the regression coefficient describing the effect of survey date on the probability of detection for forest birds on Puerto Rico surveyed before (2015) and after (2018) passage of hurricanes Irma and Maria.** Dots show the mean of the posterior samples, and lines represent the 95% credible interval. The solid horizontal line represents no effect of date on probability of detection; species falling below the line had lower detectability during surveys conducted later in the season, whereas species above the line had higher detectability during surveys conducted later in the season. For more information on species codes and names, see S1 Table.

**S3 Figure. Value of the regression coefficient describing the effect of time of day on the probability of detection for forest birds on Puerto Rico surveyed before (2015) and after (2018) passage of hurricanes Irma and Maria.** Dots show the mean of the posterior samples, and lines represent the 95% credible interval. The solid horizontal line represents no effect of time on probability of detection; species falling below the line had lower detectability during surveys conducted later in the day, whereas species above the line had higher detectability during surveys conducted later in the day. For more information on species codes and names, see S1 Table.

**S4 Figure. Relationships between the probability of detection and the probability of site occupancy among 35 species of forest birds on Puerto Rico surveyed during a winter before (2015; a) and after (2018; b) passage of hurricanes Irma and Maria.**

**S5 Figure. Probability of occupancy among forest birds surveyed on Puerto Rico during a winter before (2015) and after (2018) passage of hurricanes Irma and Maria.** Dots show the mean of the posterior samples, and lines represent the 95% credible interval. For more information on species codes and names, see S1 Table.

**S1 Table. Percentage of sites at which each species was detected and total number of sites at which a species was detected during surveys conducted at 227 points (186 surveyed in both years, 25 surveyed only in 2015, and 16 surveyed only in 2018) in forested areas across Puerto Rico during January - March 2015 and 2018.** Species in bold were included in the formal analysis of occupancy; English common names and 4-letter codes are provided as well for these species.

**S1 Appendix. JAGS code used to analyze hierarchical site-occupancy models.**

